# Structure of a dodecameric double-ferritin-fold protein from an Asgard archaeon

**DOI:** 10.64898/2026.08.25.747088

**Authors:** Alina Remeeva, Arina Anuchina, Dmitrii Dashevskii, Tikhon Kurkin, Oleg Semenov, Alexey Mishin, Stepan Osipov, Guangyi Li, Pavel Shishkin, Yaroslav Shuvaev, Anatolii Mikhailov, Elizaveta Kuznetsova, Ilia Natarov, Andrey Nikolaev, Vsevolod Sudarev, Alexey Vlasov, Valentin Borshchevskiy, Andrey Rogachev, Ivan Gushchin

## Abstract

Ferritins are ubiquitous iron homeostasis proteins found across the tree of life that form conserved 24-subunit cages with octahedral (4-3-2) symmetry. New types of ferritins and ferritin-like proteins are being continuously discovered, such as mini-bacterioferritins, which form smaller shells of 12 subunits, and double-ferritin-fold proteins, which act as ferroxidases but do not form shells. Here, we describe double-ferritin-fold proteins from Asgard archaea, dubbed dFTNs, and determine Cryo-EM structure of a representative from *Candidatus* Heimdallarchaeum endolithica. The protein forms a dodecameric shell with tetrahedral (2-3) symmetry. N-terminal (NTD) and C-terminal (CTD) domains are bridged by an ordered linker and are related by two-fold rotational pseudosymmetry. C-terminal α-helix (helix E) that forms the four-fold channel in classic ferritins is repositioned to be the helix 2 out of 5 ferritin domain α-helices in dFTN, with two such helices from NTD and two helices from CTD forming a pseudo-four-fold symmetry structural element. Four three-fold channels are formed by NTDs, and four other such channels are formed by CTDs. The overall arrangement of dFTN ferritin domains is similar to that of protomers in classic ferritin shells. Altogether, our findings expand the range of known ferritin family proteins and provide insight into Asgard archaea iron metabolism.

## Introduction

Ferritins and ferritin-like proteins are essential molecular entities for iron homeostasis across the tree of life [1,2]. They form shell-like cages that consist of either 12 or 24 subunits. Dodecamers are typical for Dps (DNA proteins from starved cells) and Dps-like ferritins; tetracosameric proteins are divided into bacterioferritins and non-heme ferritins, altogether often referred to simply as “ferritins”. Bacterioferritins (BFRs) were originally discovered in bacteria and to date were found exclusively in bacteria and archaea [3–5]. The key discerning feature of BFRs is their ability to bind heme molecules [6]. On the other hand, non-heme ferritins, which do not bind heme, are widespread in both prokaryotes and eukaryotes [7,8].

Recent studies revealed additional members of the ferritin group and the ferritin-like superfamily that confound the traditional classification. Wissink et al. discovered dodecameric mini-bacterioferritins (mBFRs) that group with BFRs in the phylogenetic tree and bind one heme-like ligand (Fe-coproporphyrin III) per two ferritin monomers [9]. Elsewhere, we reported a group of double ferritin-like proteins (DFLPs) that contain two four-helical domains; despite high similarity to ferritins, DFLPs form dimers in solution that do not assemble into shells [10].

In the present work, we describe the proteins with two ferritin domains that form oligomeric shells similar to those observed in classic ferritins and BFRs. These double ferritins (dFTNs) are restricted to Asgard archaea and TACK group [11,12], while being closely related to mini-bacterioferritins by sequence. Cryo-EM structure of a representative protein from *Candidatus* Heimdallarchaeum endolithica reveals unusual rearrangements in the ferritin domains that make dFTNs unique members of the family. Below, we describe them in detail and discuss the implications.

## Results and Discussion

### Phylogenetic analysis and prediction of oligomeric states

We began our work by examination of proteins consisting of two ferritin-like domains. For better understanding of their distribution among other ferritin-like proteins, we built a phylogenetic tree (Fig. 1), similarly to our previous study [10]. Interestingly, besides the double ferritin-like proteins (DFLPs), which were shown not to form cages [10], we found a separate cluster of proteins from Asgard archaea with two ferritin-like domains, which we dub dFTNs. N-terminal and C-terminal domains (NTDs and CTDs, respectively) of these proteins form separate clades with the same origin in the phylogenetic tree, likely indicating the emergence of dFTNs through a domain duplication event (Fig. 1). All 24 identified dFTNs (full list may be found in Supporting Information) consist of two ferritin domains, whereas 2 related sequences (UniProt IDs A0A832U0P2 and A0A7J3ACV9) appear to be truncated due to incomplete sequence assembly. Average sequence identity between NTD and CTD is 39.7% for dFTNs. Interestingly, dFTNs belong to the same clade in the phylogenetic tree as the recently discovered Mper-mBFR, which consists of a single ferritin-like domain and forms dodecameric shells [9].

**Figure 1.**
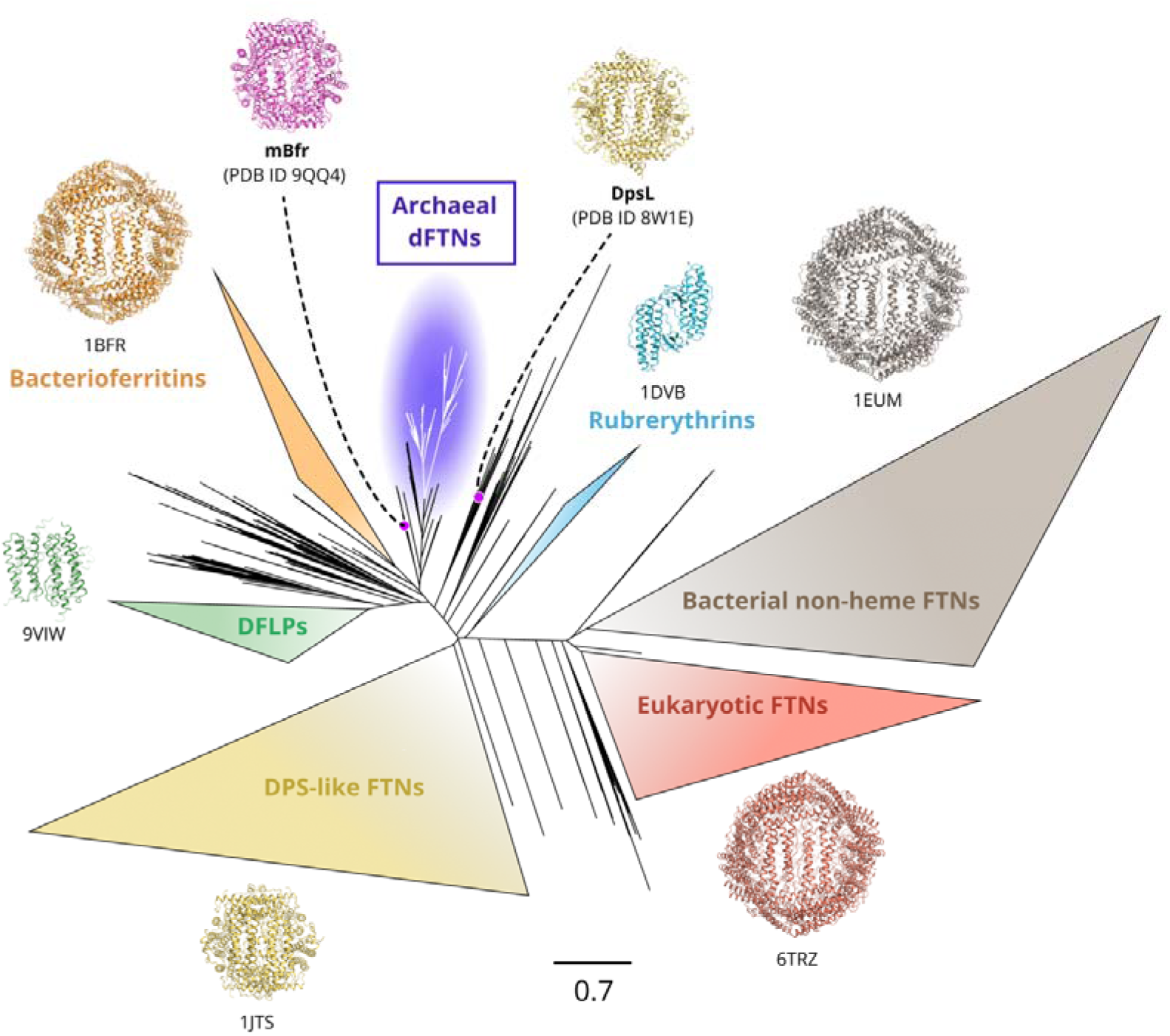
Phylogenetic tree of ferritin-like proteins. N- and C-terminal domains of newly identified dFTNs form two distinct clades and are highlighted in violet. The closest structurally characterized homolog of dFTNs is the dodecameric mini-bacterioferritin from *Candidatus* Methanoperedens carboxydivorans (mBfr, PDB ID 9QQ4).

Given our previous experience with DFLPs, which formed dimers and not ferritin-like cages, we attempted modeling various assemblies of dFTNs and related proteins using AlphaFold3 [13] (Fig. 2) prior to any experiments. We found that AlphaFold3 accurately reproduces the available experimental structures of Mper-mBFR (PDB ID 9QQ4) from *Ca.* Methanoperedens carboxydivorans and the Dps-like ferritin from *P. aeruginosa* (PDB ID 8W1E), both of which are dodecamers. Dodecameric assembly is also predicted for representatives from two other related clusters (UniProt IDs A0A0D6JTT4 and A0A015ZP02, Fig. 2). When AlphaFold3 is tasked with predicting tetracosamers of these proteins, it consistently produces two separate dodecamers (models may be found on Zenodo using the following link: https://doi.org/10.5281/zenodo.22045965). On the other hand, for the dFTN clade proteins, assembly into a BFR-like shell consisting of 24 ferritin domains is reliably predicted (Fig. 2), whereas assemblies from 6 proteins (corresponding to 12 ferritin domains) result in unusual arrangements with lower confidence (models may be found on Zenodo).

**Figure 2.**
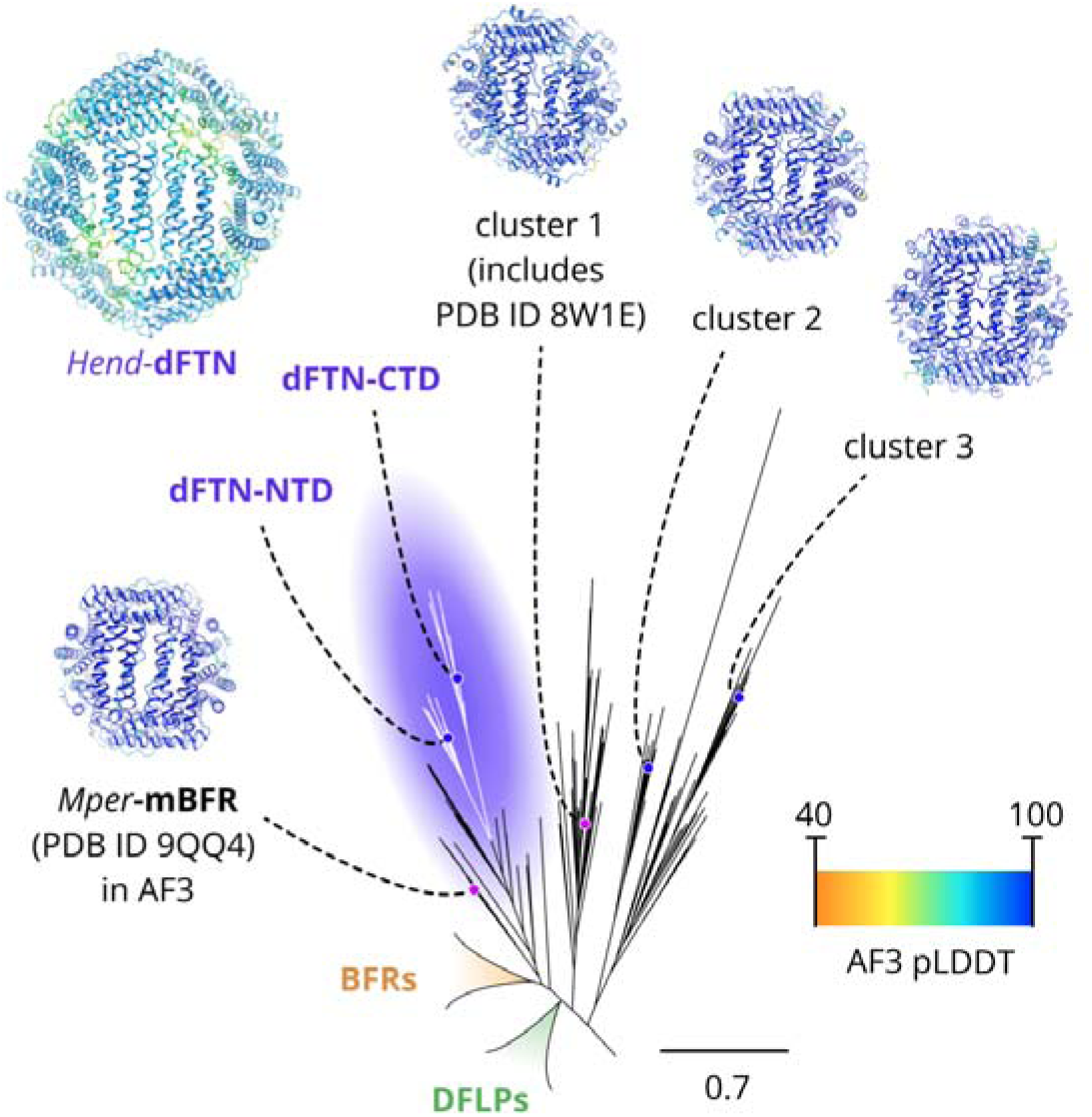
Phylogenetic context of dFTNs. AlphaFold3 modeling suggests that all related proteins form cages. The cages comprise 24 ferritin domains (12 proteins each comprising 2 domains) for dFTNs and 12 ferritin domains for proteins from adjacent clusters.

### Expression and purification of dFTNs

Encouraged by AlphaFold predictions, we selected three representative dFTNs identified in the genomes of *Ca.* Heimdallarchaeum endolithica (Hend-dFTN), *Ca.* Lokiarchaeota archaeon (Loki-dFTN) and *Ca.* Bathyarchaeota archaeon (Bathy-dFTN) for investigation *in vitro*. We attempted heterologous expression of all three proteins in *E. coli*. Loki-dFTN and Bathy-dFTN could not be purified in amounts sufficient for consequent analyses and were discarded from further study, whereas Hend-dFTN produced usable samples. In size-exclusion chromatography, Hend-dFTN eluted in a single peak corresponding to molecular weight (MW) of 400–450 kDa (Supporting Fig. 1), comparable with the predicted MW of a dodecamer of 468 kDa.

### Electron microscopy study of dFTNs

For preliminary analysis, the purified sample of Hend-dFTN was subjected to negative stain electron microscopy. Well-formed ferritin-like cages of appropriate size were easily discernible in the photographs and 2D classes (Fig. 3A). Consequently, we proceeded with single particle analysis, which produced a model with tetrahedral symmetry (Fig. 3B). The local resolution of the model varies from 2.1 Å to 3.0 Å, and the nominal resolution is 2.4 Å (Fig. 3C-E).

**Figure 3.**
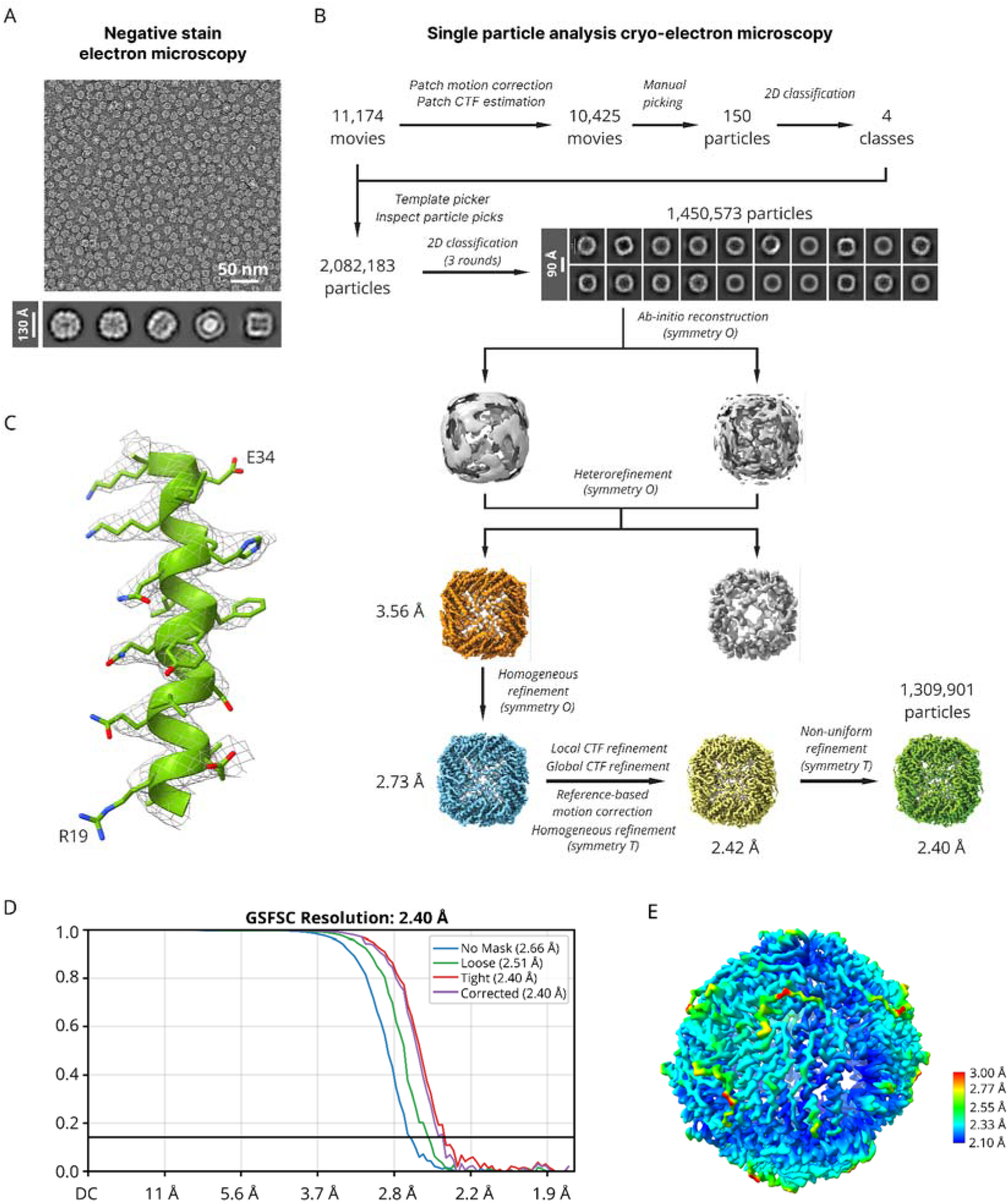
Electron microscopy study of dFTN. A) Negative stain electron microscopy. Ferritin-like cages are easily discernible in the photographs and 2D classes. B) Single particle analysis workflow. C) Final density map obtained using CryoSPARC in the region comprising residues 19-34. The map is contoured at the level of 5σ. D) Fourier-shell correlation (FSC) plots for the cryo-EM reconstruction. Resolution values at FSC level equal to 0.143 are indicated in parentheses. E) Local resolution map calculated using CryoSPARC and visualized using ChimeraX.

Overall, Hend-dFTN forms a shell highly similar to that of classic ferritins. Given that each protein in the assembly consists of two ferritin domains, only 12 copies are observed, and overall symmetry is not octahedral (4-3-2) but tetrahedral (2-3). Accordingly, 2-fold symmetry axis of classic ferritins is replaced by a pseudo 2-fold symmetry axis between the N-terminal and C-terminal domains (Fig. 4). Instead of eight identical 3-fold symmetry elements, four such elements are observed for NTD and four for CTD (Fig. 4). Finally, 4-fold symmetry axis is replaced with a pseudo 4-fold symmetry axis truly corresponding to a 2-fold symmetry axis (Fig. 4).

**Figure 4.**
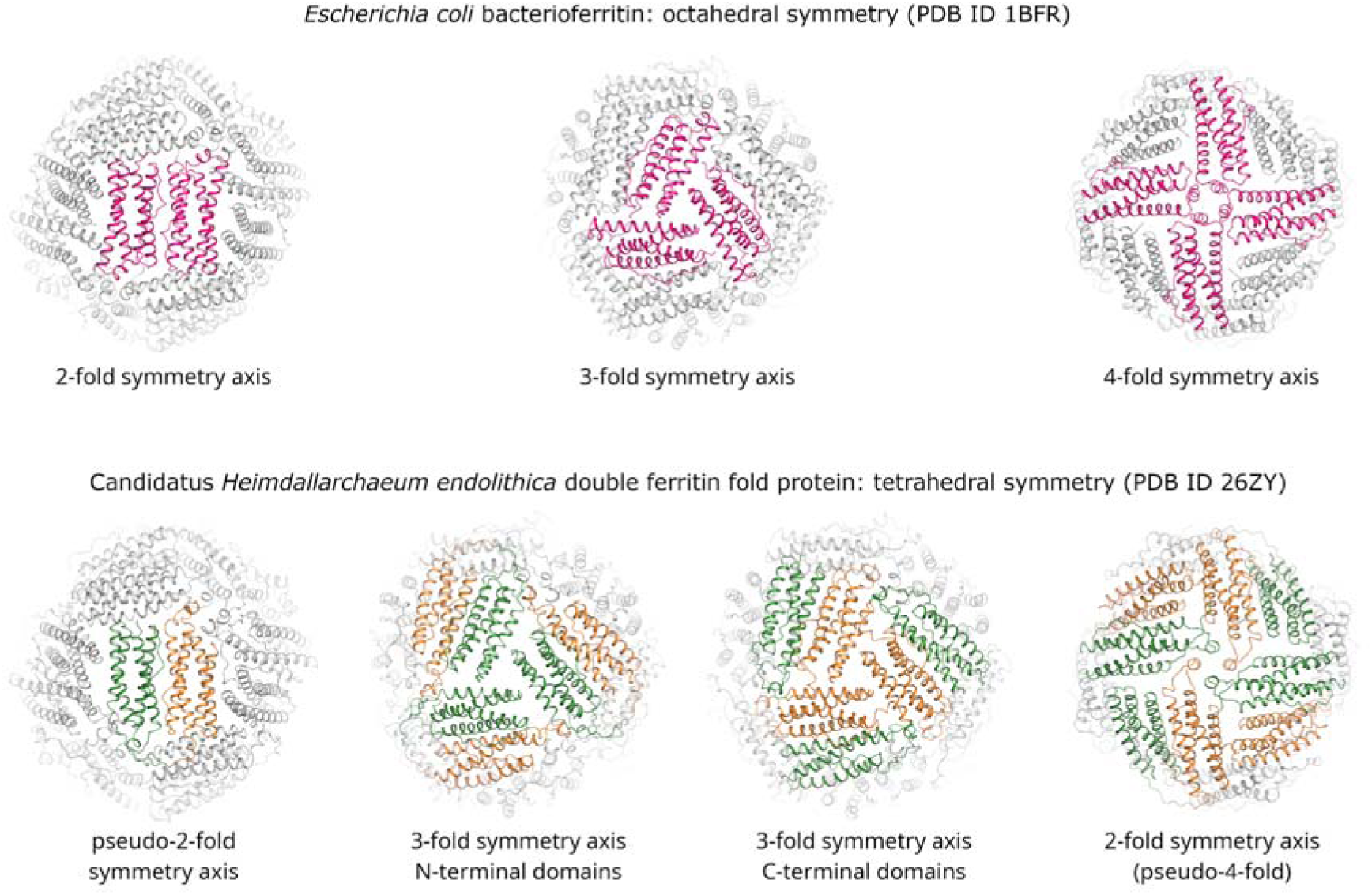
Comparison of the classic bacterioferritin assembly (top, symmetry-related protomers are shown in pink) and the dodecameric assembly of double-ferritin-fold proteins (bottom, N-terminal domains of symmetry-related protomers are shown in green and C-terminal domains are shown in orange).

NTD and CTD of Hend-dFTN are highly similar to each other, with Cα RMSD between the corresponding residues of 1.03 Å. The two domains are connected by an ordered linker (Fig. 5), unlike in DFLPs, where it is disordered [10]. Unexpectedly, the short α-helix, which forms a coiled coil at the 4-fold symmetry axis and found at the C-terminus in classic ferritins, is now found between the helices 1 and 3 in both domains of dFTN (Fig. 5); this further underscores common evolutionary origin of NTD and CTD. Sequence analysis (Fig. 6) and AlphaFold modeling shows that this short helix at this position is conserved in all 24 full-length dFTNs identified in the present study.

**Figure 5.**
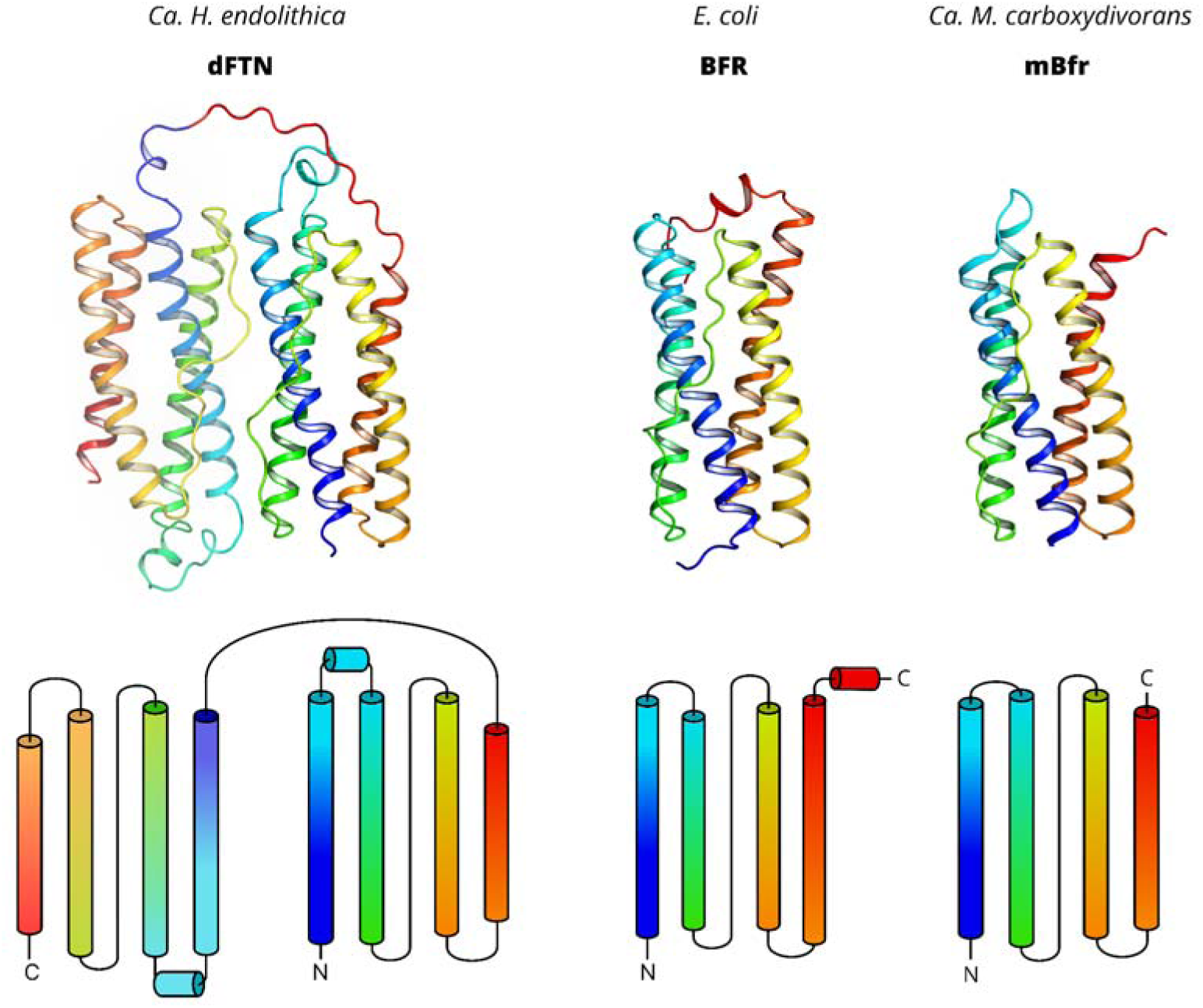
Overall folds of dFTN, BFR and mBfr. The folds differ by presence and position of a short α-helix, found after the N-terminal α-helix in dFTN and at the C-terminus in BFR, and absent in mBfr. This short helix forms a coiled coil at the 4-fold symmetry axis within the assembled shell in classic ferritins and dFTNs.

**Figure 6.**
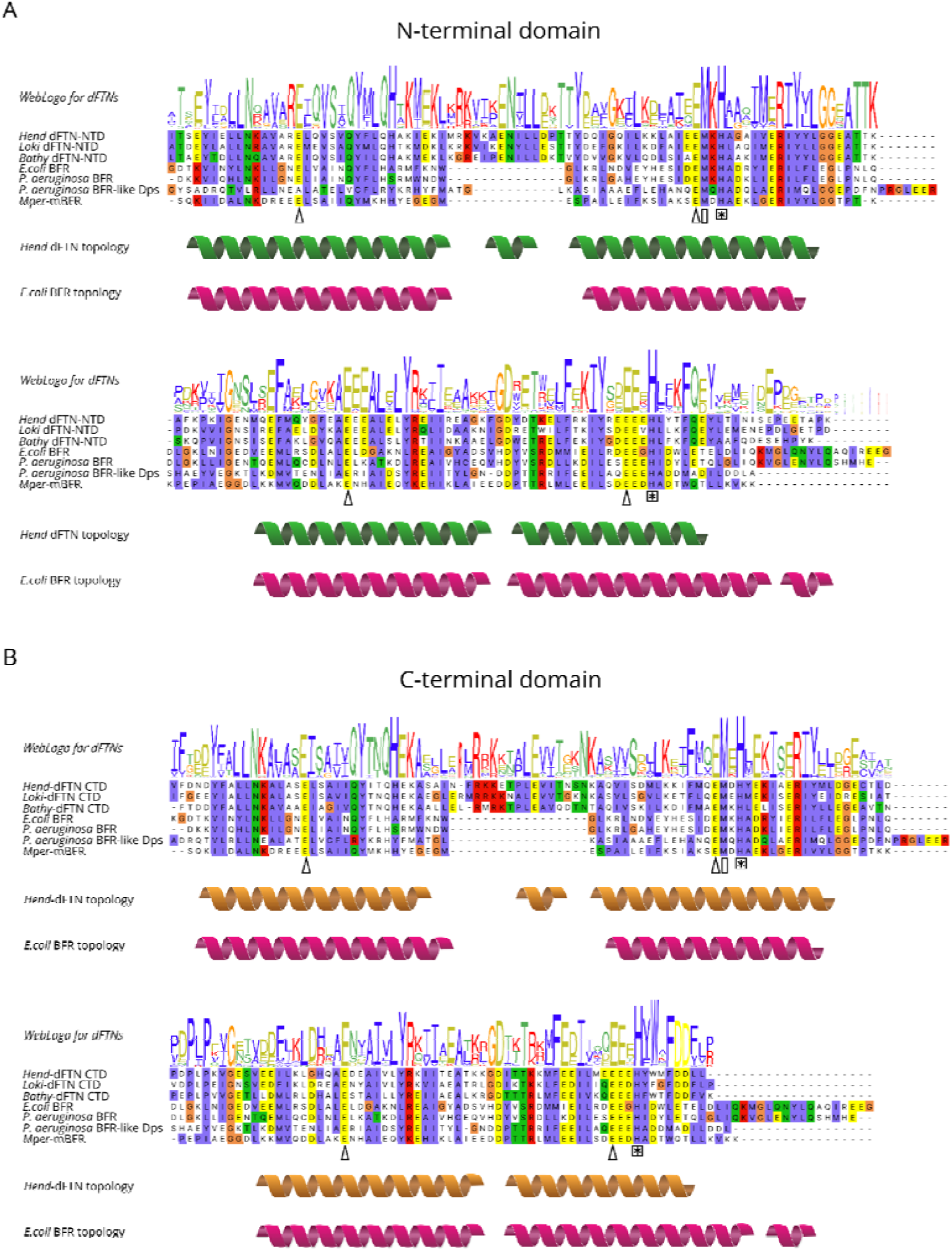
Sequence analysis of dFTNs and related proteins. WebLogo plots and sequences for the NTD (top) and CTD (bottom) of dFTNs are compared with sequences of the representative ferritin family members. Secondary structure is indicated below the sequence alignment in green, orange and magenta for NTD, CTD and classic ferritins, respectively. Conserved glutamate, histidine, and heme-ligating methionine positions are indicated with triangles, asterisks, and rectangles, respectively.

NTD and CTD domains of Hend-dFTN are related by a pseudo-2-fold symmetry axis. In classic bacterioferritins, as well as in closely related mini-bacterioferritins, corresponding interface is often occupied by a heme family molecule, whose iron is ligated by the sulfur atoms of nearby methionines (Fig. 7). Accordingly, two methionine amino acids project towards the interface in Hend-dFTN, although no ligand is observed (Fig. 7A). In the model obtained using AlphaFold 3, heme molecule is reliably placed at the interface (median pLDDT of protein atoms of 92, median pLDDT of heme atoms of 70) without the need for any large-scale conformational rearrangements (RMSD of Cα atom positions of neighboring α-helices compared to the apo-model obtained from experiment is 0.52 Å). Methionines 68 and 238, the likely iron ligands, are strongly conserved among dFTNs (Fig. 6).

**Figure 7.**
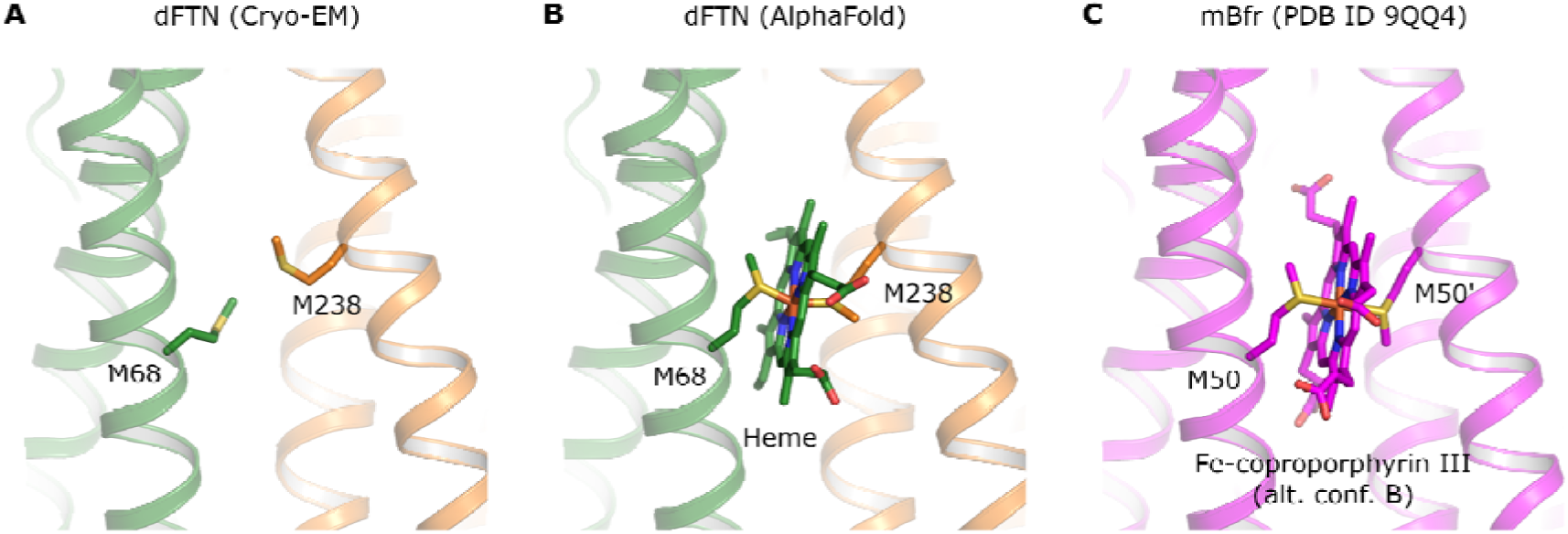
Possibility of heme binding at the NTD (green)-CTD(orange) interface of dFTN. A) Putative binding site in the Cryo-EM model of dFTN. B) AlphaFold model of dFTN bound to heme. M68 and M238 ligate the heme’s iron. RMSD of Cα atom positions of respective α-helices compared to the apo-model obtained from experiment is 0.52 Å. C) Binding site of Fe-coproporphyrin III at the interface between two mBfr protomers related by crystallographic symmetry. RMSD of Cα atom positions of respective mBfr α-helices compared to the apo-model of dFTN obtained from experiment is 0.71 Å.

3-fold symmetry centers formed by NTD and CTD differ in their physico-chemical properties. For NTD, the putative channel is lined with positively charged amino acids, with K132 and R136 occupying the centermost positions, and R117, R121 and K125 residing in close vicinity (Fig. 8A). For CTD, the putative channel is formed by two negatively charged amino acids (E307 and E308), one positively charged (K303) and one polar but neutral (T295, Fig. 8B).

**Figure 8.**
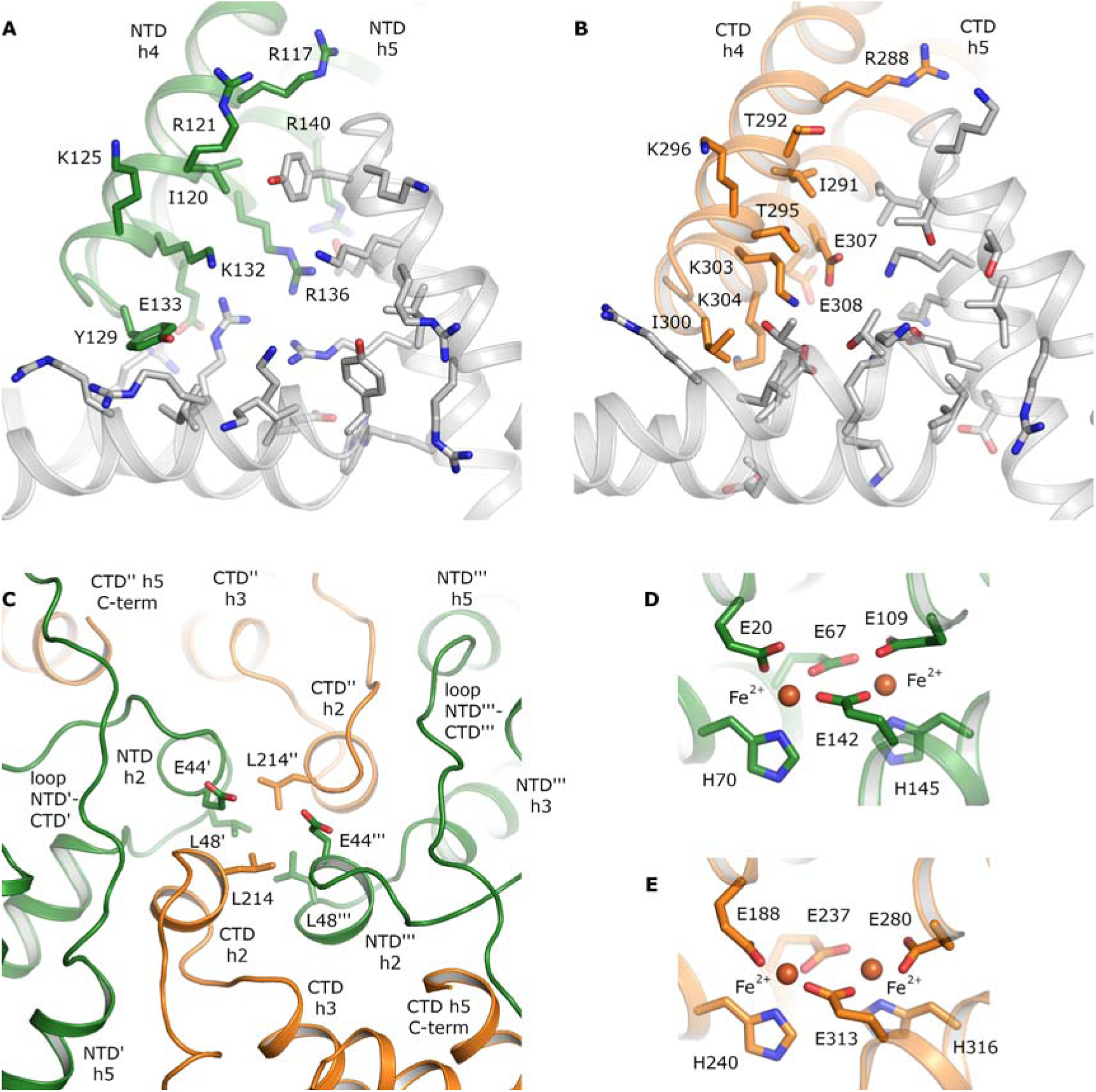
Structural elements of dodecameric ferritins. A) Three-fold channel formed by helices 4 and 5 of NTDs. One protomer is shown in green and its channel-forming residues are labeled; the two others are shown in grey. Out of 9 pore-forming residues, 6 are positively charged. B) Three-fold channel formed by helices 4 and 5 of CTDs. One protomer is shown in orange and its channel-forming residues are labeled; the two others are shown in grey. The channel has no clear charge, its constriction is lined both by positively charged (K303 and K304) and negatively charged (E307) residues. C) Pseudo-four-fold channel formed by helices 2 of NTDs and CTDs. Structural elements belonging to protomers 2, 3 and 4 are marked with 1, 2 and 3 apostrophes. D) Di-iron site in NTD. E) Di-iron site in CTD.

The pseudo-4-fold symmetry center is formed by short α-helices composed of residues 43-47 of NTD and 213-219 of CTD. At the interface of this coiled coil arrangement are hydrophobic amino acids L48 and L214, with glutamic acids E44 facing the exterior of the dFTN cage (Fig. 8C). NTD-CTD linkers are found nearby, each contacting the NTD α-helix belonging to the same protomer and CTD α-helix belonging to the adjacent protomer.

Finally, both domains have putative ferroxidase sites, each composed of 4 glutamates, 2 histidines and 2 putative Fe^2+^ ions (Fig. 8D, E). Geometry of these sites is nearly identical to that of classic ferritins. Sequence alignment of representative dFTNs, BFR, mBFR, and DpsL from *P. aeruginosa* reveals conservation of the di-iron site residues in both NTDs and CTDs of dFTNs, which suggests that the ferroxidase activity is retained within the dFTN clade (Fig. 6).

## Discussion

Iron is essential for many biochemical reactions across the tree of life, and ferritin family proteins are among the major actors in iron metabolism [1,2,14]. Accordingly, they have been thoroughly studied, and their diversity has been reliably mapped using molecular and genomic approaches [15–17]. Yet, some organisms, such as Asgard archaea, have escaped detailed studies until recently [12,18]. Whereas Asgard representatives are notoriously difficult to cultivate, availability of hundreds of their genomes allows studies of their molecular mechanisms.

Here, we used genome mining to identify unusual two-domain ferritin-like proteins in some of the Asgards. Utility of AlphaFold modeling was already demonstrated for Asgard proteins [18] and we used it here to guide the experimental work. Our analysis, initiated on May 17^th^, 2024, produced 24 full-length two-domain proteins. For *Ca.* H. endolithica, further analysis revealed another ferritin-like protein (UniProt ID A0A9Y1BQZ6) elsewhere in the genome. However, AlphaFold modeling indicates that the latter protein is highly likely to be a domain-swapped dimer rather than a cage-forming protein, structurally similar as determined using the DALI server [19] to encapsulated ferritins [20] or rubrerythrins [21,22] (Supporting Fig. 2). dFTNs have intact di-iron sites, which could oxidise iron, and form ferritin-like shells, where the iron could be stored, and thus might be the main iron storage proteins in *Ca.* H. endolithica and related organisms. At the same time, not all Asgard organisms have cage-forming ferritins of this type, as further analysis shows single ferritin domain proteins related to bacterial non-heme FTNs in genomes of other Bathyarchaeota, Lokiarchaeota, and Heimdallarchaeota.

dFTNs are closely related to mini-bacterioferritins and classic bacterioferritins. The latter were shown to bind heme or heme-like ligands that facilitate electron transfer across the shell as well as to the di-iron ferroxidase center [4,23,24]. The heme’s iron is usually ligated by the sulfur atoms of nearby methionine amino acids. Accordingly, dFTNs have conserved methionines at this position, and AlphaFold modeling indicates that they should be capable of binding heme or related compounds without any notable structural rearrangements. Yet, no coloration of Hend-dFTN samples was observed after heterologous expression in *E. coli*, and no heme-like moieties were found in the Cryo-EM densities in the corresponding region. It is possible that dFTNs bind a related but different compound, which is absent from *E. coli* (mBfr samples purified from the native source were shown to bind to Fe-coproporphyrin III), and/or the expression conditions were not suitable for any notable heme binding. Thus, it remains to be elucidated whether dFTNs bind heme-like molecules in native conditions, and what could be their role.

Structural analysis reveals that Hend-dFTN, and likely other dFTNs, form tetrahedral cages similar to octahedral cages of classic ferritins and bacterioferritins, consisting of 12 polypeptides each comprising 2 ferritin-like domains, which results in 24 domains in the complete assembly (Fig. 3). Previously, ferritin from *Archaeoglobus fulgidus* was also found to form a cage with tetrahedral symmetry consisting of 24 ferritin domains [25]. However, it consisted of 24 domains that were identical in sequence yet differed in their structural environment within the cage; on the contrary, in dFTNs, NTDs and CTDs are different in sequence yet exist in similar structural environments. The proteins that are most closely related to dFTNs, such as mBfrs (Fig. 2), form dodecamers consisting of 12 ferritin-like domains. Moreover, while having a similar overall fold, dFTNs have a notable difference from other cage-forming ferritins, as the short α-helix forming an inter-subunit coiled coil is located right after the N-terminal α-helix and not at the C-terminus. Thus, it is possible that the 24 domain assembly re-emerged during evolution of dFTNs, following loss of the short α-helix in mBfr-like proteins.

Presence of two domains in a single dFTNs polypeptide opens interesting possibilities for ferritin research and for synthetic biology. First of all, the two domains may be mutated separately, allowing one-by-one interrogation of NTD and CTD elements. Second, ferritin cages with strict 1:1 stoichiometry of two different domains may be constructed. Finally, ferritin nanoparticles are often used for presenting other proteins on their surface, e.g. in vaccine design [26–28]; dFTNs provide an opportunity of presenting only 12 and not 24 domains, allowing fusion with larger functional agents (proteins, polymers, etc) compared to traditional ferritins. These features may be useful in a wide range of biotechnological applications [29].

## Materials and Methods

### Bioinformatic analysis

The dataset containing sequences of ferritin-like proteins was assembled as described previously [10]. The primary analysis is based on the previously obtained phylogenetic tree [10], which was built using FastTree 2.0 [30], and annotated using FigTree v.1.4.4 [31]. Multiple sequence alignment was performed using MUSCLE v. 5.15.2 [32]. Domain architecture was determined using AlphaFold3 [13] and HMMER 3.3.2 [33]. Genomic environment data were obtained from GenBank [34], and corresponding IDs for genes and assemblies were obtained using the IDmapping tool in UniProt [35].

### Cloning, expression and purification

The nucleotide sequences corresponding to genes K9W46_01370 (UniProt protein ID A0A9Y1BS62, residues 3-324, dubbed Hend-dFTN), HWN80_03545 (UniProt ID A0A7K4GKX5, residues 2-326, dubbed Loki-dFTN) and GX563_01275 (UniProt ID A0A7K3ZMF0, residues 2-326, dubbed Bathy-dFTN) with N-terminal 6×His tags (full additional tag sequence MGHHHHHHSGG) were optimized for *Escherichia coli* expression, synthesized de novo and inserted into the pET-28(+) expression vector.

The proteins were expressed in *E. coli* strain BL21(DE3). Cells were cultured in shaking baffled flasks in LB medium containing 150 mg/L kanamycin. Protein expression was induced by 1 mM IPTG and continued for 5 h at 37 °C.

Harvested cells were disrupted using M-110P Lab Homogenizer (Microfluidics, USA) at 25000 psi in a lysis buffer containing 300 mM NaCl, 50 mM Tris-HCl, pH 8.0 with addition of 0.5 mM PMSF. The lysate was clarified by removal of the cell membrane fraction by ultracentrifugation at 100 000 g for 1 h at 4 °C. The supernatant was incubated with Ni-nitrilotriacetic acid (Ni-NTA) resin (Qiagen, Germany) on a rocker for 2 hours at 4 °C with addition of 20 mM imidazole. The supernatant with Ni-NTA resin was loaded on a gravity flow column and washed with a buffer containing 300 mM NaCl, 50 mM Tris-HCl, pH 8.0, supplemented by 50 mM Imidazole. The protein was eluted by a buffer containing 500 mM Imidazole, 300 mM NaCl and 50 mM Tris-HCl, pH 8.0. The eluate was subjected to a size-exclusion chromatography on a Superose® 6 Increase 10/300 GL (Cytiva, USA) column at 0.5 mL/min in a buffer containing 100 mM NaCl and 10 mM Tris-HCl, pH 8.0.

### Negative stain electron microscopy

Formvar/Carbon-coated 300-mesh Cu-grids were glow-discharged using PELCO easiGlow instrument for 15 s at 15 mA current with negative polarity. 3.5 μl of sample was applied to the glow-discharged grids for 1 min, followed by negative staining with 1 % w/v uranyl acetate solution for 1 min. After subsequent blotting, the grids were allowed to dry for 24 h. The prepared grids were then imaged on JEOL JEM-2100 transmission electron microscope operating at an accelerating voltage of 200 kV. The microscope was equipped with a DE-20 direct electron detector (Direct Electron, USA). Image acquisition was performed using SerialEM software, with pixel size being 1.4 Å. A total amount of acquired micrographs was 245. Obtained micrographs were then processed using CryoSPARC v.4.6.2 software [36] to perform 2D-classification of the particles.

### Cryo-EM sample preparation and data collection

A 3 μl aliquot sample of 0.2 mg/ml concentration was applied onto Quantifoil holey carbon supported copper grids (R1.2/1.3, 200 mesh) freshly glow-discharged for 40 s using Solarus II Plasma Cleaner (Gatan, USA). Cryo-EM grids were prepared using a Vitrobot Mark IV System (Thermo Fisher Scientific, USA) operated at 100% humidity and 4°C. Grids were blotted for 4.5 s using 595-grade ashless filter paper (Thermo Fisher Scientific, USA) with blot force 0 and immediately plunge-frozen into liquid ethane. Grids were imaged using a 300 keV Titan Krios G1 (Thermo Fisher Scientific, USA) microscope. Images were recorded using a Gatan K3 direct electron detector in super-resolution mode. The pixel size was set to 0.87 A. Exposure was set to 15 e^-^/A^2^/sec and 40 frames were collected in total, with an overall dose of 50 e^-^/A^2^.

### Image processing and map calculation

All movies were imported and analyzed in CryoSPARC v.4.6.2 and subjected to motion correction, and the contrast transfer function (CTF) parameters were estimated. We initially used CTF-corrected micrographs to manually pick 150 particles with the box size set to 270 pixels. 2D classification was performed and the most populated classes were selected for template-based particle picking. A total of 2,082,220 particles were extracted from 10,425 micrographs as a result of auto-picking and subsequent picks inspection. After a series of 2D classifications, 1,450,573 particles were used for *ab initio* reconstruction with octahedral O symmetry and 2 classes. After heterogeneous refinement, the class with resolution of 3.56 Å (FSC = 0.143) was chosen for the subsequent homogeneous refinement with O symmetry that improved resolution to 2.73 Å (1,312,278 particles). Global and local CTF refinement, reference-based motion correction and one more round of homogeneous refinement with tetrahedral T symmetry were applied and improved resolution to 2.42 Å (1,309,901 particles). The result of non-uniform refinement with resolution of 2.40 Å corresponds to the final map. The processing strategy is illustrated in Fig. 3 and statistics are presented in Table 1.

**Table 1.** Cryo-EM data collection, refinement, and validation statistics.

|  |  |
| --- | --- |
|  | **dFTN protomer** |
| PDB ID | 26ZY |
| EMDB ID | EMD-81006 |
|  | **Data Collection and Processing** |
| Magnification | 81 000 |
| Voltage (kV) | 300 |
| Electron exposure dose (e<sup>-</sup>/Å<sup>2</sup>) | 50 |
| Pixel size (Å) | 0.87 |
| Symmetry | T |
| № of movies | 10425 |
| № of initial particle images | 2082220 |
| № of final particle images | 1309901 |
| Map resolution (Å) | 2.40 |
|  | **Refinement** |
| Model resolution (Å) | 2.40 |
|  | **Model composition** |
| № of non-hydrogen atoms | 2650 |
| № of protein residues | 320 |
| № of water molecules | 3 |
| № of iron atoms | 4 |
|  | **B factors (Å<sup>2</sup>)** |
| Protein | 101.45 |
| Iron | 65.77 |
| Water | 60.37 |
|  | **R.M.S. deviations** |
| Bond lengths (Å) | 0.011 |
| Bond angles (°) | 2.157 |
|  | **Validation** |
| MolProbity score | 1.39 |
| Clash score | 7.02 |
| Rotamer outliers (%) | 0.70 |
|  | **Ramachandran plot** |
| Favored (%) | 98.43 |
| Allowed (%) | 1.57 |
| Outliers (%) | 0.00 |

### Model building and refinement

The model of a protomer generated using AlphaFold3 [13] was fitted using rigid body docking into the map with ChimeraX-1.10.1 [37]. As the cryo-EM map reconstruction possesses tetrahedral T symmetry, all symmetry elements of the apoferritin shell are recapitulated if one protomer (the asymmetric unit) is suitably positioned in the tetrahedral point-group symmetry T. Therefore, all subsequent refinement steps were performed with the asymmetric unit and the final model was generated by applying the T symmetry for the asymmetric unit. Сoordinates of the apoferritin protomer (asymmetric unit) were refined in Phenix 2.0-5936 [38] using phenix.real_space_refine. Then CCP-EM Doppio Refmac Servalcat [39] was used for further refinement and generation of difference Fourier maps, which served as guides for manual refinement. After automatic refinement, the protomer model was manually inspected and side-chain fits were refined in COOT v0.9.8.96 [40].

## Data availability

Density maps were deposited into the Electron Microscopy Data Bank under the accession code EMD-81006 and atomic coordinates were deposited into the Protein Data Bank under the accession code 26ZY. AlphaFold models were deposited to Zenodo (https://doi.org/10.5281/zenodo.22045965).

## Supporting information

Supporting information file is available that includes Supporting Figures 1-2 and Supporting Data with a list of identified dFTN proteins.

## Contributor Roles

Conceptualization: I.G. Funding acquisition: I.G., V.B., A.M. Investigation: A.R., A.A., D.D., T.K., O.S., A.M., S.O., G.L., P.S., Y.S., A.M., E.K., I.N., A.N., V.S., A.V., V.B., A.R., I.G. Project administration: A.R., A.A., I.G. Writing – original draft: A.R., A.A., D.D., I.G. Writing – review & editing: A.R., A.A., D.D., T.K., O.S., A.M., S.O., G.L., P.S., Y.S., A.M., E.K., I.N., A.N., V.S., A.V., V.B., A.R., I.G.

## Acknowledgements

Bioinformatic and structural analyses as well as sample preparation were supported by the Ministry of Science and Higher Education of the Russian Federation (agreement 075-03-2026-305, project FSMG-2025-0003). Negative stain EM experiments and optimization procedures were supported by the Russian Science Foundation (project No. 22-74-10036-П). Cryo-EM data collection was supported by the Ministry of Science and Higher Education of the Russian Federation (agreement 075-15-2025-512). Cryo-EM data treatment and model building were supported by the Ministry of Science and Higher Education of the Russian Federation (agreement # 075-03-2026-305, project FSMG-2024-0012). We thank the staff members of Electron Microscopy System (https://cstr.cn/31129.02.NFPS.EMAS) at the National Facility for Protein Science in Shanghai for providing technical support and assistance in data collection.

