## Supporting Information for "Structure of a dodecameric double-ferritin-fold protein from an Asgard archaeon"

Supporting Figures 1-2

Supporting Data

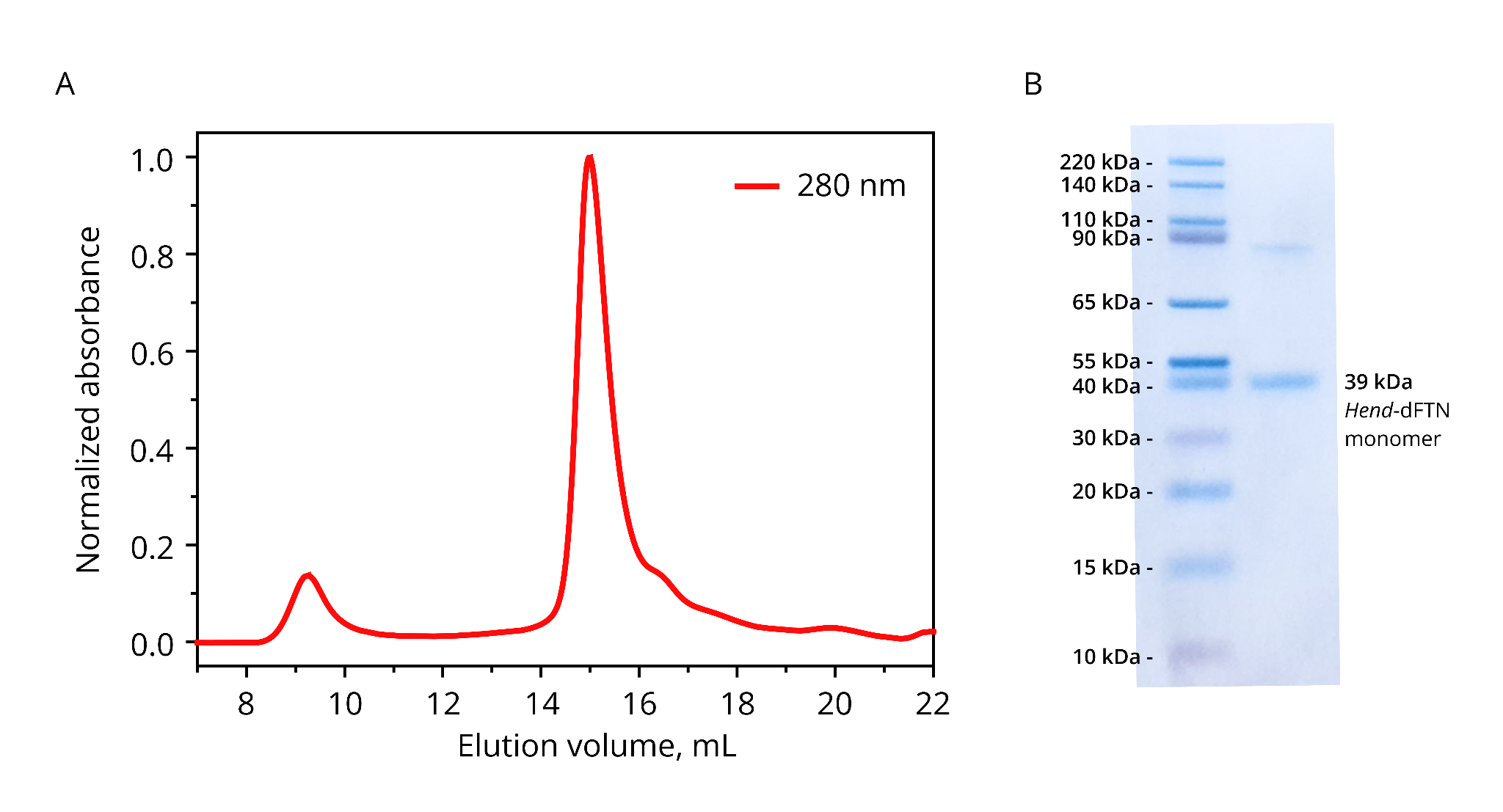

**Supporting Figure 1.** Purification of dFTN from *Ca.* Heimdallarchaeum endolithica. A) SEC profile of the dFTN sample following Ni-NTA chromatography. Chromatography was performed using a Superose® 6 Increase 10/300 GL column. The peak is observed at 15 mL, which corresponds to dodecameric assembly of the protein with resulting MW of 400–450 kDa. B) SDS-PAGE analysis of the peak fraction in denaturing conditions. The second band with MW of ~80 kDa presumably corresponds to dFTN dimers.

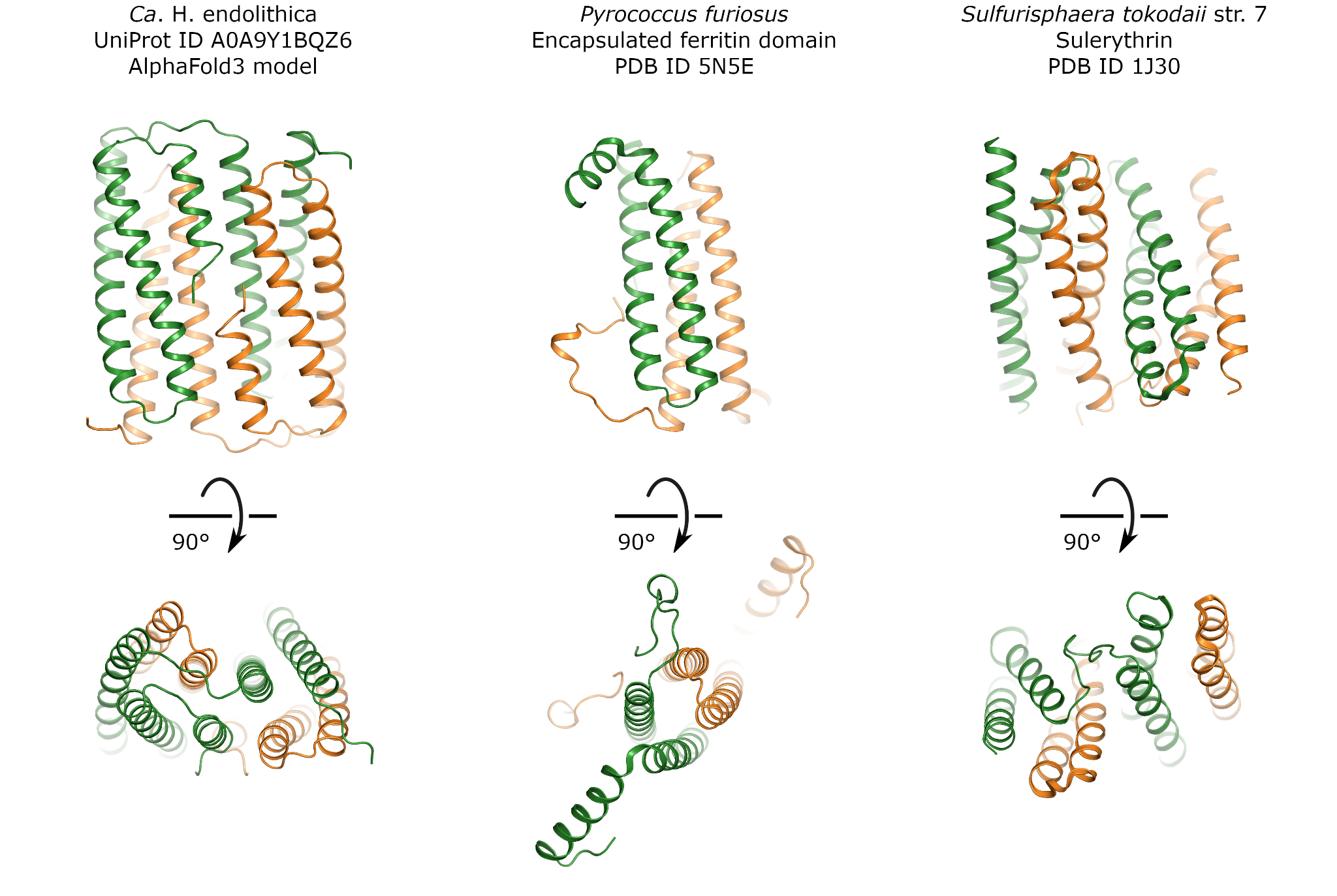

**Supporting Figure 2.** Comparison of a ferritin-like protein from *Ca.* H. endolithica (UniProt ID A0A9Y1BQZ6) with the closest structurally characterized ferritin family proteins. In each case, one protomer is shown in green and another one is in orange. A) AlphaFold3 model of ferritin-like protein UniProt ID A0A9Y1BQZ6 from *Ca.* H. endolithica. Average pLDDT is 86.1 and median pLDDT is 90.3. The protein is predicted with high confidence to be a dimer with domain swapping present (exchange of two α-helices between the two domains). Models may be found on Zenodo using the following link: https://doi.org/10.5281/zenodo.22045965. B) Structurally related encapsulated ferritin from *Pyrococcus furiosus* (PDB ID 5N5E, DALI server RMSD of 4.5 Å, sequence identity of 14%). The protein is a decamer; only two protomers forming a single ferritin-like domain are shown. C) Structurally related sulerythrin (*Sulfolobus* rubrerythrin-like protein) from *Sulfurisphaera tokodaii* str. 7 (PDB ID 1J30, DALI server RMSD of 4.8 Å, sequence identity of 10%). The protein is a dimer with domain swapping present, yet the relative arrangement of the two ferritin-like domains is different from that predicted for the *Ca.* H. endolithica protein.

**Supporting Data 1.** dFTN proteins identified in this study.

| UniProt ID | Organism | Number of amino acids | Amino acid sequence |
| --- | --- | --- | --- |
| A0A9Y1BS62 | *Candidatus* Heimdallarchaeum endolithica | 324 | MSVKITSEYIELLNKAVARELQVSVQYFLQHAKIEKIMRKVKAENILLDPTTYDQIGQILKKLAIEEMKHAGAIVERIYYLGGEATTKAFKPKIGENMQEFMQYGFEAEEEALELYREIIREAGKFGDYDTKELFRKIYREEEEHLYTFQEYLTINISEPEETAPKSEWREVFDNDYFALLNKALASELSAIIQYITQHEKASAINFRKKETPLEVITNSNKAQVISDMLKKIFMQEMDHYEKIAERIYMLDGECTLDPDPLPKVGESVEEILKLGHQAEDEAIVLYRKIITEATKKGDITTKKMFEEILMEEEEHYWMFDDYL |
| A0A8T5TCZ4 | *Promethearchaeota* archaeon | 326 | MIIKATNEYLALLNQAVAREIQVSAQYMLQHTKIDKLKRKVIKENYLLESTTYDEFGKILKDLAIEEMKHLAQIMQRIYLLGGEATTKPDRVIIGNSMKEFAELNLKAEEEALELYRQLIEAAKNIGDRETWALFTKIYSDEEVHLLKFQDYVDMEAEPDLGETPASEWRSIFGDEYIALLNKALASEISAVVQYTTQHEKAEGLERMRRKKNALEVVTGKNKASVVSGVLKEIFLQEMDHMEKISERIYEIDRESIATLDPLPEIGNSVDEFIKLDREAENYAIVLYRKVIAEATRLGDIKTEKLFEDIIQEEEEHYWAFDDFLP |
| A0A8T5TTC7 | *Promethearchaeota* archaeon | 326 | MIIKATDEYLALLNRAAAREIQVSAQYMLQHTKMEKLKRKVIKENYLLETTTYDEFGEILKDLAKEEMKHLAQIMGRIYLLGGEATTKPDRVIIGNSLREFAELDVKAEEEALELYRILIEAAKDIGDRETWVLFTKIYSDEEEHFLKFQDYVEMESEPDLGKTPDSEWRSIFGEEYVTLLNKALASEISAVVQYTNQHEKAEGLERMRRKKNALEVVTGKNKESIVSGILKETFMQEMEHMEKISDRIHEIDRDSIATVDPLPEIGNSIDEFIKLDRDAENYAIVLYRKIITEATRLGDIKTRKLFEDIIVEEEAHYWNFDDFLP |
| A0A0F9HIJ7 | marine sediment metagenome | 329 | MSIKFEKPTEEYIELLNRSVAREIQVSTQYMIQHTKMEKLLRKVIKENILLDTTTYEEFGKILKEFAVQEMKHLGKIIERIYILGGEATTKPDKIIIGDSLREFAELGVKAEEEALKLYHKIIEEAKNIGDRETWLLFSKIYSDEEEHLLKFQDYSEIEDEPDLGETPESDWRSIFKSDYIALLNQALSSEISAIIQYTTQHEKAEGLERMRRKKAALEVVTGKNKASVVSGILQEIFIQEMEHMEKIAERIYEINREALIQVDPLPEIGNTVDDFIKLDRNAENYAIVLYRRVIEKATELGDIKTKVMFEDIIQQEEEHYWKFDDFLP |
| A0A7K4GKX5 | *Promethearchaeota* archaeon (“*Candidatus* Lokiarchaeota” archaeon) | 326 | MIVKATDEYLALLNRAVAREMEVSAQYMLQHTKMDKLKRKVIKENYLLESTTYDEFGKILKDFAIEEMKHLAQIMERIYLLGGEATTKPDKVVIGNSIREFAELDYKAEEEALELYRQLIDAAKNIGDRETWILFTKIYSDEEVHLLKFQEYLEMENEPDLGETPDSEWRSIFGEEYIALLNKALASEISAVIQYTNQHEKAEGLERMRRKKNALEVVTGKNKASVLSGVLKETFLQEMEHMEKISERIYEIDRESIATVDPLPEIGNSVEDFIKLDREAENYAIVLYRKVIAEATRLGDIKTKKLFEDIIIQEEDHYFGFDDFLP |
| A0A8T5RD93 | *Promethearchaeota* archaeon | 300 | partial sequence  YMLQHTKIEKLRRKVIKENYLLEDTTYDEFGEILKDLAKEEMKHLAQIMARIYLLGGEATTKPDRVIIGNSLREFAELDVKAEEEALELYRILIEAAKDIGDRETWVLFTKIYSDEEEHFLKFQDYVEMENEPDLGETPDSEWRSIFGEEYVALLNKALASEISAVVQYTNQHEKAEGLERMRRKKNALEVVTGKNKASVVSGILKETFLQEMEHMEKISDRIYEIDRESIATIDPLPEIGNSVEDFIKLDREAENYAIVLYRKVIAEATRLGDIKTRKLFEDIIQDEETHYWAFDDFLP |
| A0A7K4HFR2 | *Promethearchaeota* archaeon | 326 | MIVKATNEYLALLNSAAAREIQVSAQYMLQHTKMEKLRRKVIKENYLLEDTTYDEFGEILKDLAKEEMKHLAQIMARIYLLGGEATTKPDRVIIGNSLREFAELDVKAEEEALELYRIIIEAAKDIGDLETWVLFTKIYSDEEKHFLKFQDYVEMESEPDLGGTPDSEWRSIFGEEYVTLLNKALASEISAVVQYTNQHEKAEGLERMRRKKNALEVVTGKNKASVVSGILKETFLQEMEHMEKISDRIYEIDRESIATVDPLPEIGNSVDEFIKLDREAENYAIVLYRKIIAEATRLGDIKTRKLFEDIIQDEESHYWAFDDFLP |
| A0A8T5UQW0 | *Promethearchaeota* archaeon | 326 | MIVKVTDEYLGLLNRAAAREIQVSAQYMLQHTKMEKLKRKVIKENYLLETTTYDEFGEILKDLAKEEMKHLAQIMGRIYLLGGEATTKPDTVIIGNSLREFAELDVKAEEEALELYRILIEAAKDIGDRETWVLFTKIYSDEEEHFLKFQDYVEMENEPDLGETPDSEWRSIFGEDYVTLLNKALASEISAVVQYTNQHEKAEGLERMRRKKNALEVATGKNKASVVSGILKETFLQEMEHMEKISDRIHEIDRDSIATVDPLPEIGNTVDDFIKLDREAENYAIVLYRKVIAEATRLGDIKTRKLFEDIIQDEETHYWAFDDFLP |
| A0A7K3YLN1 | *Thermoproteota* archaeon | 326 | MAVKITTDYIDMLNQTVEHEFRVSIQYFLQHAKMEKLKKRANPENILLDKTTYDAVGKVFYDISVSEMKHAADIMERIYYLGGQATTKSSTPVIGNSISEFAKLGIETEKEALTLYRKIIPAAREAGDWQARKLFERIYSEEEEHLFKFQEYANFQDEKNESNEATKPEWQKIFTDDYFALLNKAVAAEITGIIQYTNQHEKAALLGNRLKNTPLEAIQNTNIAGVTGNILKDIFLVEMNHLEMISERIYLLGGEVTANPNPLPVVGDTVFDMLKLDHSLESMTISLYREIIAEALKRGDTTTRLIFEEIAKQEEEHFWTFDDFVK |
| A0A7K4CV13 | *Promethearchaeota* archaeon | 325 | MVEKPSAELVALLNAGVARELQVSAQYILQHTKMEKLLRKVRAENILLETTTYEALGKLLKTMAIEEMKHAGTIMERLYYLGAEATTKAAPVKIGKNLKEFMSHGFEAEAEALELYMKTIKLAEGEGDMETGEIFRKIYSDELKHYHSFEERLKLDISEPEGPKDIESKHTSVYTSEYFSLLNKAVAAEIGAIVQYTNQHEKASKIALRSLETPMEVIGEKNKAAVVSEMLKKFARQEMSHLDKIAERIHLLGGDVVTTPDPLPKIGETVDDFLRNGKEGEDYAIVLYRQIIAKAIEIGDITTRKVFEGITDEEDGHYWAFDDYF |
| A0A9Y1BL59 | *Candidatus* Heimdallarchaeum aukensis | 324 | MSVKITSEYIELLNKAVARELQVSVQYFLQHAKIEKIMRKVKAENILLDSTTYDQIGQILKKLAIEEMKHAGSIVERIYYLGGKATTKAFKPKIGENMQEFMKYGFEAEEEALELYREIIREAGKLGDYDTKELFRKIYREEEEHLYTFQEYLTINISEPGETAPKSEWRKVFDDEYFALLNKALASELSAIIQYITQHEKASAINLRKKETPLEVITNTNKAQVISNMLKKIFMQEMEHYEKISERIYMLDGECTLDPDPLPKVGESVEDFLKLGHQAEDEAIVLYRKIIEEATKKGDITTKKMFEEILMEEEEHYWMFDDYL |
| A0A7J3R159 | “*Candidatus* Bathyarchaeota” archaeon | 325 | MAVEVSSEYIDLLNQAVARELQVSIQYMLQHAKMEKLIRRTLSENILLDKTTYDAVGKFLREFAIQEMKHAAAVMERIYYLGGTATTKANRVNVGNSISEFARNGVKAEEEALALYRKIIDSTGRVGDVETRELFEKIYGEEEKHLFKFQEYVNVQDETGESQMSLSDWRKIFSEDYFTLLNKAVAAEISAIVQYTNQHEKASLLALRTKNTPLEVITEANKAKVVSDMLKPIFMVEMEHLEKITERIYLLEGEAVSEPEPIPKVGETAEEFLRLDHEAENYAIVLYRKIIEESLKRGDTTTRRLFEDIVMQEEGHYWQFDDFLR |
| A0A7J2TBE2 | “*Candidatus* Bathyarchaeota” archaeon | 326 | MAVQPTSEYFDLLNQAVSREIQVSIQYILQHAKMEKLMRKVIPENMLLDKTTYEAVGKFLKEIAIQEMKHAADIMERIYYLGGSATTKSNKPVIGNSLSEFAKLGAKAEEEALVLYRKIIEAAKALGDVETWKMFEKIYSQEEQHLFKFQEYVNMKDEPEDAEAQPVSEWRKIFTDDYFALLNRAVAAEISAIIQYTNQHEKASLLALREKVSPLEVVTESNKAKVISDLLKKIFMVEMEHLEKISERIYLLEGECTVTPDPIPQVGETADDFVKLDHEAENIAIVLYRQIIAEALKRGDTTTRRMFEDIVLQEEEHYWAFDDFLR |
| A0A7K3ZMF0 | “*Candidatus* Bathyarchaeota” archaeon | 326 | MVIKLTAEYTDLLNQAVAREIQVSIQYILQHAKMEKLKGREIPENILLDKTVYDVVGKVLQDLSIAEMKHAAKIMERIYYLGGEATTKSKQPVIGNSISEFAKLGVQAEEEALSLYRTIINKAAELGDWETRELFEKIYGDEEEHLFKFQEYAAFQDESEHPYKVPMPEWRKIFTDDYFALLNKAVAAEIAGIVQYTNQHEKAALLELRMRKTPLEAVQDTNTAQIVSKILKDIFMAEMKHLELISERIYLLEGEAVTNPEPLPVVGETLLDMLRLDHALESTAILLYREIIAEALKRGDTTTRLMFEEIVKQEEEHFWTFDDFVK |
| A0A1F5DVM6 | “*Candidatus* Bathyarchaeota” archaeon RBG_13_52_12 | 325 | MAVKMTSDYVDLLNEAVARELQVSIQYMLQHTKMEKLIRKVIPENILLDKTTYEAVGKFLKEISIQEMKHAAAIMERIYYLGGQATTKSKKPVVGGSLSEFAKLGVEAEEEALILYRRIIDESRKVGDYESHELFEKIYGEEEGHLFKFQEYVKVRDESEGDSGETSEWRKIYTEDYFALLNKAVASEISAIVQYTNQHEKAALLSLRMKETPLEVITEKNKTKAISDLLKGIFMQEMEHLEKISERIYLLEGEATVNPEPLPKVGDTADDFLRLDHKAENDAIVLYRKIIEEAMKRGDTLTRRMFEDIVIQEEGHYWKFDDYLR |
| A0A938YQA8 | “*Candidatus* Bathyarchaeota” archaeon | 324 | MAVKVTSEYIGLLNQAVSREIQVSIQYMLQHTKMEKLMRKAIPENILLDKTTYDAVGAFLKEIAIEEMKHAADIMERIYLLGGSATTKADKPVVGGSLSEFARLGVAAEAEALELYRKIIKASREVGDRTTRQLFQKIYKAEEEHLLKFQEYVDLKDEPEDAESAVSEWRKIFTDDYFALLNKAVASEISAIIQYTNQHEKASLLALRKKVSALEVVTESNKAKIVSDLLKTVFMQEMDHLEKISERIYLLGGECTVVPDPIPQVGADPKDFIALDKDAENTAIMLYRQIIAEALKRGDTTTRNMFEEIIKQEEEHYWSFDDYQ |
| A0A8T5U7C2 | *Promethearchaeota* archaeon | 326 | MIVKATDEYLALLNRAAAREIQVSAQYMLQHTKIEKLRRKVIKENYLLEDTTYDEFGEILKDLAKEEMKHLAQIMARIYLLGGEATTKPDTVIIGNSLREFAELDVKAEEEALELYRILIEAAKDIGDRETWVLFTKIYSDEEKHFLKFQDYIEMENEPDLGETPDSEWRSIFGEEYVALLNKALASEISAVVQYTNQHEKAEGLERMRRKKNALEVVTGKNKASVVSGILKETFLQEMEHMEKISDRIYEIDRESIATVNPLPEIGNLVEDFIKLDREAENYAIVLYRKIIAEATRLGDIKTRKLFEDIIQDEETHYWNFDDFLP |
| A0A7J3JFN0 | “*Candidatus* Bathyarchaeota” archaeon | 326 | MSVKITTEYIDMLNKAVEREIGVSLQYILQHAKMEKLMRKTLPENILLDKTTYDAVGKFLKDIAIQEMKHAATIMERIYYLGGQATTKSSKVTVGDSLSEFAKLGVKAEEEALVLYRQIIETATKMGDWETHEVFEKIYGEEEGHLFKFQEYTKFQDEKDEPSKVPLPEWRKIYTDDYFALLNKAVAAEITGIVQYTNQHEKAAFLELRRKNTPLETITETNKADVVSKLLKGVFMQEMEHLEKISERIYLLEGEAVAKPDPLPVVGETAQDFLILDHALESSAITLYRQIIAEALKRGDTTTRRLFEDIVMQEEEHFWSFDDFIR |
| A0A8T5U4Z5 | *Promethearchaeota* archaeon | 326 | MIIKATNEYFALLNRAAAREIQVSAQYMLQHTKMEKLKRKVIKENYLLESTTYDEFGKILKDLAIEEMKHLAQIMERIYLLGGEATTKPDKVIIGNSLREFAELDYKAEEEALELYRQLIEAAKDIGDRETWVLFSKIYSDEEEHLLKFQDYLEMENEPDLGETPDSEWRSIFGEAYITLLNKALASEISAVVQYTTQHEKAEGLERLRRKKNALEVVTGKNKASIVSGVLKETFLQEMEHIEKISERIYEIDRESIAKIDPLPEIGNSVDDFIKLDRKAENYAIVLYRKVITEAIRLGDIKTKKLFEDIIQQEEEHYWAFDDFLP |
| A0A7J4NYW8 | “*Candidatus* Bathyarchaeota” archaeon | 325 | MAVKTTPKYIDLLNEAVEREIQVSIQYLLQHGKMEKLIRRTLPENILLDKTTYEAVGKFLREIGIQEMKHAAAIMERIYYLGGKATTKAKKPVIGGSLSEFAKLGVGAEEEALTLYRKIIDEARKAGDYETHELFEKIYGEEEVHLFKFQEYTKFKDEPQDSDVKPSEWRRAYTDDYFALLNKAVASEIMAIIQYTNQHEKAALLKLRMKETPLETITEKNKTQVVSDLLKPIFMQEMEHLEKISERIYLLEGEAVAMPDPLPKVGETVDEFLHLDHEAENDAILLYRRIIDEAMKRGDTLTRRMFEDIVIQEEEHYWKFDDFIR |
| A0A1Q9MSI5 | *Promethearchaeota* archaeon (strain CR_4) | 327 | MMIVKTTTPEYFDLLNKGVSRELQVSVQYMLQHSKMQKLLRRVIPENMLLDKTTYEALAKVLQEFAIQEMKHAGAIMERIYLLGGQATTKADKVKIGDSLREFGTLDVKAEEEALDLYRKVIEMAAKLGDWETREMFEKIYGDEEKHLIRFQEFTEVADEPDFGTTAGSEFETVFKDDWIAMLNKALASEISAIIQYTNQHEKANVEQNRRRKTALEVVTDKTKPISISDLTRSIAMDEMKHMEAIAERIYEIKKECVAAVDPLPDVGDTADNWIMNNRVAENSAITFYRQIISKALEIGDVKTRKMFEDIIVQEEDHYWKFDEWVP |
| A0A7C3TB92 | “*Candidatus* Bathyarchaeota” archaeon | 326 | MAVKATSDYFELLNKAVSREIQVSIQYMLQHGKMEKLMRKVIPENILLDKTTYEAVGKFLREFAIMEMKHAASVMERIYYLGGSATTKSSKPNIGNSLSEFAKYGLKAEEEALVLYRNIIDAARKVEDWETRELFEKIYSDEEKHLFKFQEYVNIQDEPEGSEQAPLSEWRRIFTSDYFDLLNKAVAAEISAIIQYTNQHEKASLLALREKNTALEVVTEKNKAKVVSELLIPIFKAEMEHLEKISDRIYLLEGEATTEPDPLPQIGETVDDFLKLDHEAENYALVLYRKIIDEAMKRGDSKTRRMFEDIIGQEEDHYWTFDDFLR |
| A0A7J2YSV3 | “*Candidatus* Bathyarchaeota” archaeon | 326 | MAVNITTEYIDLLNQAVEREFQVSIQYFLQHSKMEKIKKREIPENLLLDKTVYDAMGKILEDTAIMEMKHAAAIMERIYYLGGAATTKAKKITVGDSISEFAKLDLKAEEEALILYRQIIETAAKMGDWETRELFEKIYGEEEDHLFKFQEYTEFQDEKEEPYKVPMPEWRKIFTDDYFALLNKAVAAEITGILQYTNQHEKAALLEFRRKSTQLEGVQGTNKAEVVSKILKDIFMVEMKHFEMITERIYLLDGECVTEPDPKPVVGATVQDALVLDHALESSAIFLYRQIIAEALKRGDTTTRLMFEEIVKQEEEHFWTFDDFVR |
| A0A842R0E4 | “*Candidatus* Bathyarchaeota” archaeon | 325 | MAIKPTSELLKMLNQGVARELQVSVQYMLQHFKMERILRKVRKENILLEGTTYESLGGILKQMAIEEMKHLADIMERIYYLGGKATTKSDKPQIGENLKDFMEFGYKAEEEALELYRKVITEAEKIGDWETAEMFKEIYRQEEEHLYTFEEYLTVDITEPEGPEDVPTDSVKIYTDDYFELLNKAVAAEISAIVQYTNQHEKASKLALRKKEKPMEVIKSKNKASVISDLLKEVFMKEMDHLEMISERIYLLGGEAVYNPYPLPVIGETVDDFLRLDKKAEDYAIVLYRQIVAEATKLGDTVTKRMFESILEDEDQHYWMFDDYF |
| A0A832U0P2 | “*Candidatus* Bathyarchaeota” archaeon | 272 | partial sequence  AVGKFLKEFAIQEMKHAASVMERIYYLGGEATTKGNKPSIGNSLSEFARNGVKAEEEALVLYRKIIEAAGKIGDWETREVFEKIYGEEESHLFKFQEYSKMQDEAEGPPAVPLSEWRKIFTDDYFALLNKAVQAEISAIIQYTNQHEKASLQALRERNTALEVVTESNKPSVVSKLLKGIFMQEMEHLEKISERIYLLEGEAVFTPDPIPKVGSNADDFLKLDHEAENIAILLYRKIVAEALKIGDTKTRRLFEDIVMQEEEHYWTFDDYVR |
| A0A7J3ACV9 | “*Candidatus* Bathyarchaeota” archaeon | 233 | partial sequence  IGNSLSEFAKLGAKAEEEALVLYRKIIEAAKALGDVETWKMFEKIYSQEEQHLFKFQEYVNMKDEPEDAEAQPVSEWRKIFTDDYFALLNRAVAAEISAIIQYTNQHEKASLLALREKVSPLEVVTESNKAKVISDLLKKIFMVEMEHLEKISERIYLLEGECTVTPDPIPQVGETADDFVKLDHEAENIAIVLYRQIIAEALKRGDTTTRRMFEDIVLQEEEHYWAFDDFLR |
